# The Soil Food Web Ontology: aligning trophic groups, processes, resources, and dietary traits to support food-web research

**DOI:** 10.1101/2023.02.03.526812

**Authors:** Nicolas Le Guillarme, Mickael Hedde, Anton M. Potapov, Carlos A. Martínez-Muñoz, Matty P. Berg, Maria J.I. Briones, Irene Calderón-Sanou, Florine Degrune, Karin Hohberg, Camille Martinez-Almoyna, Benjamin Pey, David J. Russell, Wilfried Thuiller

## Abstract

Although soil ecology has benefited from recent advances in describing the functional and trophic traits of soil organisms, data reuse for large-scale soil food-web reconstructions still faces challenges. These obstacles include: (1) most data on the trophic interactions and feeding behaviour of soil organisms being scattered across disparate repositories, without well-established standard for describing and structuring trophic datasets; (2) the existence of various competing terms, rather than consensus, to delineate feeding-related concepts such as diets, trophic groups, feeding processes, resource types, leading to ambiguities that hinder meaningful data integration from different studies; (3) considerable divergence in the trophic classification of numerous soil organisms, or even the lack of such classifications, leading to discrepancies in the resolution of reconstructed food webs and complicating the reuse and comparison of food-web models within synthetic studies. To address these issues, we introduce the Soil Food Web Ontology, a novel formal conceptual framework designed to foster agreement on the trophic ecology of soil organisms. This ontology represents a collaborative and ongoing endeavour aimed at establishing consensus and formal definitions for the array of concepts relevant to soil trophic ecology. Its primary objective is to enhance the accessibility, interpretation, combination, reuse, and automated processing of trophic data. By harmonising the terminology and fundamental principles of soil trophic ecology, we anticipate that the Soil Food Web Ontology will improve knowledge management within the field. It will help soil ecologists to better harness existing information regarding the feeding behaviours of soil organisms, facilitate more robust trophic classifications, streamline the reconstruction of soil food webs, and ultimately render food-web research more inclusive, reusable and reproducible.

## Introduction

Trophic interactions mediate the ecosystem processes provided by soils, including carbon sequestration, nutrient cycling, pest and pathogen regulation. These soil ecological processes also support aboveground biodiversity, soil health, ecosystem services and ultimately ecosystem resilience and stability [1]. Modelling energy and nutrient transfers between soil organisms through accurate reconstructions of soil food webs provides a better understanding of the relationships between multitrophic assemblages of soil organisms and ecosystem functioning [2]. However, comprehensive soil food-web studies remain relatively rare due to the high demand for trophic data and the absence of easy-to-use soil food-web modelling tools [3].

The methodological toolbox for studying trophic interactions and answering the ‘who-eats-whom-and-what?’ question in soil has expanded considerably over the last two decades [4, 5], resulting in a rapid accumulation of data on the feeding behaviours of soil organisms. Simultaneously, there is a growing call for a more holistic and integrative soil ecological research [6]. Yet, the integration of data remains challenging, primarily due to the prevailing practice of data collection by individual teams for their specific projects, often without considering data standardisation. This results in the dispersion of data across various publications, databases, or general-purpose file hosting services [7]. In addition, soil trophic ecology encompasses a diverse range of concepts, including diets, food resources, trophic interactions, guilds, trophic groups, and feeding processes [4]. These terms are often subject to multiple interpretations, have variable meanings in different contexts, or lack widely accepted and unambiguous definitions [8, 9]. This terminological ambiguity and the lack of data and metadata standardisation hamper the efficient reuse and synthesis of published datasets, as it makes it difficult to find, interpret and integrate relevant data into broader-scale studies [10].

In the last decade, some progress has been made towards the creation of standard terminologies for describing, sharing and facilitating the aggregation of biodiversity data, e.g. organismal occurrence [11] or trait [12, 13]. Still, standardised vocabularies for harmonising trophic datasets in soil ecology are scarce. A notable exception is the Thesaurus for Soil Invertebrate Trait-based Approaches (T-SITA) [9], which provides agreed-upon definitions for approximately 100 traits and ecological preferences for soil invertebrates, including about thirty terms describing feeding behaviours (diets and foraging strategies). T-SITA has been successfully used as the unifying terminology in the Biological and Ecological Traits of Soil Invertebrates (BETSI) database, which compiles soil invertebrate trait data from multiple sources [14]. However, T-SITA’s coverage of soil food-web concepts is actually very limited. Furthermore, T-SITA only represents hierarchical (broader/narrower), equivalence (synonym) and associative (related) relationships between concepts. As a result, T-SITA is unable to represent complex relationships between trophic concepts, such as those linking diets to resources consumed, in a formal (i.e. standardised and machine-interpretable) way.

In this paper, we build on previous efforts to harmonise and standardise terms related to the trophic ecology of soil organisms to develop the Soil Food Web Ontology (SFWO), an **<u>OWL ontology</u>** that provides a formal representation of the terminology and concepts in the field of soil trophic ecology. Ontologies encode the knowledge in a domain (e.g. soil ecology) as sets of terms and concepts interrelated using mathematical logic [10]. This enables a precise expression of the meaning of concepts that is machine processable, ultimately allowing the use of **<u>automated reasoning</u>** to check the logical consistency of the ontology and generate new knowledge, e.g. inferring new relationships between concepts.

Ontologies have been very successful in structuring knowledge in many areas of biological and biomedical research [15–17], and are increasingly being used to describe ecological data [18–20]. The proposed SFWO aims to become the standard vocabulary and formal model of reference for soil food-web research, with its terminology of over 1000 terms (including around 500 synonyms) for describing trophic interactions, diets, food resources, trophic processes and trophic groups. Most of these terms are provided with both textual and logical definitions that were agreed upon by soil ecology experts, thus helping to resolve certain terminological ambiguities. The SFWO terminology can be used to describe the content of trophic datasets which improves the discoverability and interpretability of data and facilitates data integration to the point where it can be partially automated [21]. The rich SFWO terminological and relational information can also be used to markup soil ecology text documents with ontological concepts, thus enabling ontology-based literature searches [22, 23] and the automatic extraction of relevant information from text to create or expand literature-based trophic datasets [24, 25]. Last but not least, the formal foundations of the SFWO, grounded in mathematical logic, make it possible to derive additional knowledge from explicitly stated factual data e.g. to deduce the trophic classification of an organism e.g., (phytosaprophage) from knowledge of its food resources (e.g., decaying plant material) — using automated reasoning.

This article provides an overview of the current version of the Soil Food Web Ontology (v2023-08-23) and how it can contribute to soil food-web research. Our goal is to encourage the community of (soil) ecologists to adopt the SFWO to facilitate interoperability and reuse of published datasets, produce more standardised data in the future, and to engage experts in the field to contribute to its continuous development.

## Materials and Methods

### The Soil Food Web Ontology

The backbone of the SFWO^1^ is a set of **<u>classes</u>**, describing the main concepts in the domain of soil trophic ecology, arranged in a hierarchical manner. A class can have subclasses that represent more specific concepts that inherit all the characteristics of the parent concept. For example, the class *insectivore* would generally be defined as a subclass of the class *carnivore*. In addition to classes describing concepts, an ontology also defines a set of **<u>properties</u>**which describe how concepts are related to other concepts. Each class and property in an ontology has a unique identifier (called an IRI, for Internationalized Resource Identifier), and is assigned a label (in *italics* in the rest of the article), i.e. a string — ideally, the preferred name for the class/property — that serves as a readable name for users. A class or property may also have a number of exact, related, broad or narrow synonyms.

Figure 1 depicts the basic core structure of the SFWO, which consists of four main branches containing classes describing concepts related to organisms (*organismal entity*), food resources (*food resource*), feeding-related functional traits (*trophic quality*), and trophic processes (*trophic process*). Classes in the SFWO are interconnected using five core object properties, namely *trophically interacts with, member of*, *has input*, *has quality*, and *capable of*. Following best practices of ontology development, the SFWO reuses classes and properties from existing ontologies as much as possible [26]. For instance, all the properties used in the SFWO are imported from the Relations Ontology^2^ (RO). Also, the SFWO uses the Basic Formal Ontology (BFO) [27] as a **<u>top-level ontology</u>**. This facilitates interoperability between ontologies that are built on these common foundations.

**Figure 1.**
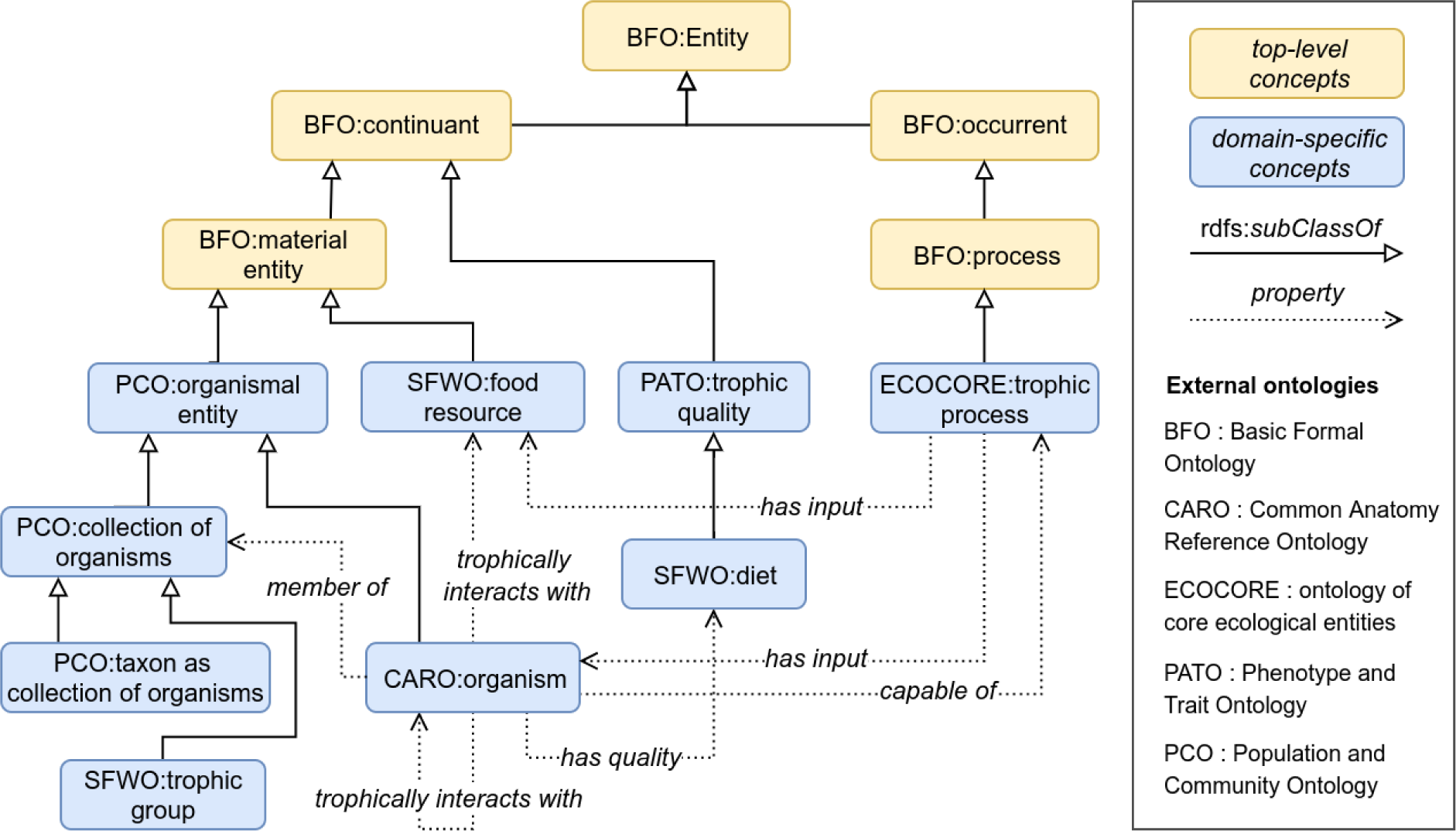
The core classes (rounded rectangles) and properties (arrows) of the Soil Food Web Ontology (SFWO). A solid arrow between two classes denotes a subClassOf relation, e.g. trophic group is a subclass of collection of organisms. Dotted arrows represent semantic relationships between classes. An organism (e.g. a detritivore) is a member of a taxonomic class (e.g. Oribatida) and trophically interacts with other organisms and/or food resources (e.g. detritus) as part of some trophic process (e.g. detritivory). Depending on the resources it consumes, an organism possesses some trophic qualities (e.g. detritivorous) and is member of one or several trophic groups (e.g. detritivorous oribatid mites).

#### Trophic quality

The SFWO imports the class *trophic quality* from the Phenotype And Trait Ontology (PATO) as the root class of a branch of the ontology dedicated to feeding-related trait classes, e.g. dietary traits, including concepts such as *bacterivorous, xylophagous, saprotrophic*, etc. It is important to emphasise that the term *quality* as used in the PATO refers to phenotypic qualities, i.e. observable properties, attributes, characteristics, traits. A *trophic quality* should therefore be interpreted as a feeding-related characteristic of an organism, not as the nutritional quality of a food resource. The SFWO introduces a new class *diet* as a subclass of *trophic quality*, defined as “a trophic quality inhering in a bearer by virtue of the bearer’s disposition to acquire nutrition by consuming certain food resources”. The *diet* class is further specialised into a number of subclasses based on the type of food resource that is consumed as part of that specific diet. For instance, the subclass *detritivorous* is defined as “a trophic quality inhering in an organism by virtue of the organism’s disposition to acquire nutrition by consuming detritus”. The class *detritivorous* is further broken down into more specialised subclasses. These include the class *phytosaprophagous,* which is defined as “a trophic quality inhering in an organism by virtue of the organism’s disposition to acquire nutrition by consuming dead or decaying plant material”. Where possible, dietary classes in the SFWO are mapped to the corresponding terms in T-SITA to make both conceptual models interoperable.

#### Organismal entity

The class *organismal entity* is imported from the Population and Community Ontology (PCO) [19], together with two of its subclasses, namely *organism* (describing a single organism) and *collection of organisms* (describing a group of two or more organisms). In the SFWO, subclasses of *organism* include (1) classes used to describe an organism according to how it feeds, e.g. *heterotroph, bacterivore, grazer*, and (2) Non-Taxonomic terms for Groups of Organism (NTGOs) [28], e.g. *microorganism*, *microalgae*, *protist*. Subclasses of *collection of organisms* include (3) *taxon as collection of organisms* (imported from PCO) for describing taxonomic classes, e.g. *Lumbricidae*, *Insecta*, *Fungi*, and (4) a new class *trophic group* as the root class of a hierarchical classification of soil-associated consumers into hybrid trophic groups.

1. *Nutrition-related organismal classes.* The SFWO provides a classification of organisms according to their feeding behaviour, including their mode of nutrition, food type, and food acquisition strategy. Much of this classification is imported from the ECOCORE^3^ ontology of core ecological entities, to which we have added many concepts related to soil food webs that were missing. The mode of nutrition refers to an organism’s source of carbon/energy/electron, e.g. *autotroph*, *heterotroph, phototroph, chemotroph, chemoheterotroph,* etc. Among these coarse-grained classes, the *heterotroph* branch is by far the most important and most developed branch of the SFWO. This branch contains a hierarchy of classes describing organisms according to their food type — e.g. *nematophage*, *coprophage*, *xylophage* — or food acquisition strategy, e.g. *scraper*, *sucker*, *shredder, suspension feeder*. The first set of classes describes an organism in terms of the specific foods (i.e. food resources or taxa) it is specialised to eat, e.g. *a springtail feeder* preys on members of the taxon *Collembola*, a *coprophage* feeds on *faeces*, etc. These classes are axiomatized to make them machine-interpretable and amenable to automatic classification using a **<u>reasoner</u>**. More specifically, we provide logical definitions of these classes in the form of equivalent class axioms, e.g. <u>equivalentTo</u>(*nematophage, organism* <u>and</u> *eats* <u>some</u> (*member of* <u>some</u> *Nematoda*)), which states that the class *nematophage* is logically equivalent to the class of all organisms that feed on (*eats*) nematodes. We also provide logical definitions that link organism classes to the corresponding diet classes, e.g. <u>equivalentTo</u>(*nematophage, organism* <u>and</u> *has quality* <u>some</u> *nematophagous*), which states that the class *nematophage* is logically equivalent to the class of all organisms that possess the trophic quality (i.e. feeding-related functional trait) *nematophagous*. Such logical definitions of classes are intended for computers, as opposed to annotations like textual definitions of classes which are primarily aimed at humans. For example, they allow a reasoner to automatically infer the dietary traits of organisms based on their trophic interactions (consumer-resource relationships).
2. *Non-Taxonomic terms for Groups of Organism (NTGOs).* Some terms for organisms commonly found in soil food-web literature cannot be associated with a single taxonomic group [28]. This is the case for example of “microalgae” which is an informal term for a large and diverse group of organisms that includes cyanobacteria and single-celled photosynthetic protists. Microalgae are the primary food source of algivorous organisms in soil. The SFWO defines classes for a number of such NTGOs which are needed to axiomatize diet classes, e.g. *microalgivore*, *protistivore, microbivore*.
3. *Taxon as collection of organisms*. All taxonomic classes are imported from the NCBITaxon ontology^4^ as subclasses of the *taxon as collection of organisms* class. The NCBITaxon ontology is an automatic translation of the NCBI taxonomy database [29] into OWL. It is used as a taxonomic reference by most ontologies maintained as part of the Open Biomedical Ontologies (OBO) Foundry [30] initiative. Other taxonomy ontologies include the BIOfid taxonomy ontology [31] (derived from the GBIF backbone) and the EUdaphobase Taxonomy Ontology^5^. We plan to add support for these and other taxonomic references (e.g. the GBIF Backbone Taxonomy) in future versions of the SFWO. To remain as compact as possible, the SFWO imports a taxonomic class (and any superclasses in its taxonomic classification path) only if that class is needed to **<u>axiomatize</u>**another class, i.e. to provide a class with a logical definition that is primarily directed at machines. For instance, the class *Nematoda* is imported from the NCBITaxon ontology to be used as part of the logical definition of the class *nematophage*.
4. *Trophic group.* To deal with the immense diversity of soil organisms when studying multitrophic assemblages, soil biologists commonly resort to classifying belowground biodiversity into “feeding guilds” of organisms that feed on the same resources, or into “trophic groups” of organisms that both feed on the same resources and have the same consumers [8]. This allows the construction of tractable food-web models across the whole spectrum of soil organisms, thus simplifying food web analysis [32]. Historically, soil organism classification into trophic groups was impeded by a persistent lack of consistency in definitions and terminology and by the lack of an overarching framework for classifying all soil biota based on their feeding preferences [8]. In the absence of such a framework, researchers had to resort to user- or clade-specific definitions of trophic groups, which led to heterogeneities in the resolution of food-web models and limited our ability to draw generic conclusions across studies [33]. Recently, Potapov et al. [4] developed an integrative classification of soil consumers from protists to vertebrates. This classification uses a hybrid taxonomic-ecological approach similar to traditional soil food-web models, e.g. Hunt et al. [34], including groups such as Oribatida-microbivores, Nematoda-fungivores. This classification provides harmonised definitions of trophic groups across different taxonomic groups, and defines a consistent aggregation strategy for food-web reconstruction. The SFWO provides a formalisation of Potapov et al.’s classification of soil-associated consumers (animals and protists) under the root class *trophic group –* defined according to Hedde et al. [8]. This hybrid classification distinguishes trophic groups individually within each taxonomic group, which makes it possible to incorporate taxon-dependent trait information in the trophic group definitions. A trophic group in the SFWO is therefore a combination of a taxon, e.g. *Nematoda, Rotifera*, and a (possibly empty) list of traits, including feeding-related traits such as *algivorous*, *detritivorous*, *bacterivorous*, etc. For example, the SFWO defines the trophic group class *Rotifera.bacterivores* using the following equivalence axiom: equivalentTo(*Rotifera.bacterivores, trophic group* and *has member* only *((member of* some *Rotifera)* and *bacterivore))*, according to which *Rotifera.bacterivores* is a trophic group composed exclusively of organisms that are both bacterivores and rotifers. This restriction allows a reasoner to infer that an organism that is a member of the trophic group *Rotifera.bacterivores* is necessarily an instance of the class *bacterivore* and a member of the taxonomic class *Rotifera*. The SFWO also provides additional subclass axioms, e.g. subClassOf((*member of* some *Rotifera*) and *bacterivore*, *member of* value *Rotifera.bacterivores*) which allows a reasoner to automatically classify an organism that is both a member of the class *Rotifera* and an instance of the class *bacterivore* as a member of the trophic group *Rotifera.bacterivores*. This makes the SFWO a valuable tool for automatically assigning organisms to trophic groups, facilitating standardised food-web reconstruction. In addition to Potapov et al.’s classification of soil-associated animals and protists and adopting the same modelling approach, the SFWO provides a formalisation of the functional classification for fungi and fungus-like organisms used by the FungalTraits database [35] and inherited from FUNGuild [36].

#### Food resource

The SFWO defines a *food resource* as “a material entity consumed to provide energetic and nutritional support for an organism”. In other words, any entity that is the object of a trophic interaction is *de facto* a food resource. In the SFWO, as in most trophic datasets, this may be a taxon, (e.g. *Aphididae* as the primary food source of aphidophagous organisms), a NTGO (e.g. *protist* as the primary food source of protistivores), organism parts (e.g. *pollen*, *blood*, *mycelium*), and other organic or inorganic materials (e.g. detritus, soil organic matter). Many of these concepts are imported from specialised ontologies, e.g. the Plant Ontology (PO) [17] for plant parts, the Fungal Anatomy Ontology^6^ (FAO) for fungal parts, the Environment Ontology (ENVO) [37] for environmental material. We decided to create a convenience class *food resource* in order to bring these very different entities, which are commonly considered as trophic resources in the literature or in food-web reconstructions, under a common root concept. This could be useful for text mining applications — for example, to extract mentions of food resources from the literature as part of a trophic interaction extraction pipeline — and for querying trophic databases.

#### Trophic processes

Mirroring the hierarchy of nutrition-related organismal classes, the SFWO provides a hierarchy of terms referring to feeding processes under the root term *trophic process*. For the sake of completeness, all the organismal classes have their processual counterparts. For example, *coprophagy* is the trophic process displayed by a *coprophage*. Although mostly useful for building a comprehensive list of terms related to trophic ecology, these process classes are also used to axiomatize the corresponding organism classes, e.g.

equivalentTo(*nematophage, organism* and *capable of* some *nematophagy*).

This logical definition uses the object property *capable of*, which describes a relationship between an entity (e.g. an organism) and a process that the entity has the ability to carry out. These axiomatisations lead to a complete mapping between trophic groups, diets, food resources and trophic processes through the intermediary of organism classes.

#### Trophic interactions

Trophic interactions are relational concepts linking a consumer to a resource. They are represented in the SFWO using object properties from the Relations Ontology. In particular, the SFWO imports an ecological subset of the RO that comprises many terms for representing biotic interactions, including *symbiotically interacts with* (and its subproperties, e.g. *parasite of*, *host of*) and *trophically interacts with* (and its subproperties, e.g. *eats*, *preys on*). Note that *parasite of* is not considered to be a trophic interaction but a symbiotic interaction in the RO, this latter being defined as “a biotic interaction in which the two organisms live together in more or less intimate association”. In its current version, the RO is missing terms to represent symbiotrophic interactions in which one organism acquires nutrients through a symbiotic relationship with another organism, e.g. trophic parasitism, mycorrhizal associations, etc. The object property *symbiotrophically interacts with* was created in the SFWO as a temporary workaround, and a new term request has been submitted to the RO GitHub issue tracker^7^.

### Building the Soil Food Web Ontology

The development of the SFWO started with a small team of experts in soil invertebrate taxonomy and ecology providing a list of over 100 terms related to primary food groups, diets and food resources of soil-associated consumers, their definitions and relationships between diets and food resources. A subset of these core terms was manually aligned with classes from existing ontologies using Ontobee’s search engine [38]. These classes were used as “seeds’’ for module extraction, a popular strategy for ontology reuse that avoids the overheads involved in importing complete ontologies [39]. Modules were merged to form the backbone of the SFWO. New classes for the terms that could not be imported from existing ontologies were created manually using Protégé^8^, a free and open-source ontology editor. The ontology development workflow is managed and automated using the Ontology Development Kit [40]. The ontology is under version control in a GitHub repository^9^ which is also used to track changes and issues, and manage releases.

The development of the SFWO has highlighted a number of terminological ambiguities currently present in the domain of soil ecology. The following examples are representative of the challenges in harmonising the terminology in a scientific domain and modelling these concepts as part of a formal representation of the domain. In our decisions we tried to adhere to the most widely accepted use of terms, rather than creating a *de-novo* (perfectly consistent) system. The project’s GitHub issue tracker allows anyone to view and contribute to discussions on these and other issues related to the ambiguous use of terms in soil food-web research.

Among the most evident issues was the terminological diversity of diets and trophic processes in soil food webs. For example, organisms feeding on fungi are named fungivores (e.g. [41]), mycophages, or mycotrophs in different literature sources (e.g. [42]). After checking the literature and expert advice, we uniformly used *-vore* and *-phage* as synonyms for organisms that feed on specific food resource by consuming the whole or part of it (*herbivore* = *phytophage*, *bacterivore* = *bacteriophage*); *-troph* is commonly used as a broader term depicting any type of energy/nutrient acquisition from specific food resource, including for instance symbiotic exchanges (symbiotrophs, such as mycorrhizal fungi) or extracellular degradation and intake of organic matter (microbial decomposition process, implemented by saprotrophs). At present, the SFWO lacks the expressiveness to represent the different modes of digestion (internal/external). As a temporary workaround, we have introduced a taxonomic constraint in some class logical definitions to distinguish between *-vore/-phage* (animals) and *-troph* (microbes). Another (open) terminological issue deals with the term algae which refers to a large and diverse group of both macro and microscopic photosynthetic organisms. In soil science, however, the term algae is used as a convenient synonym for microalgae, as most algae in soil are unicellular. In this context, it is problematic to reuse the class *algivore* from ECOCORE, as this ontology considers *algivore* to be a subclass of *herbivore*, whereas soil ecologists generally consider (micro)algivores to be microbivores, i.e. feeding on microorganisms, such as bacteria, fungi, and/or protists – the latter comprising many soil microalgae [43]. Instead, we decided to create a new class *microalgae* as a subclass of *microorganism* (with *algae* being a broad synonym for *microalgae*) and *microalgivore* as a subclass of *microbivore*. This better reflects the common understanding of these terms in soil ecology, as well as the fact that soil algae are selectively consumed mainly by bacterial, fungal, and protist feeders, rather than by vascular plant herbivores, e.g. root-feeding nematodes and curculionid larvae [4]. However, this decision is suboptimal and we probably need a term to depict all trophic processes related to primary production [28].

We also discussed the scope and classification of *detritus*. Moore et al. [44] defined detritus based on Swift et al. [45] as “… any form of non-living organic matter, including different types of plant tissue (e.g. leaf litter, dead wood, aquatic macrophytes, algae), animal tissue (carrion), dead microbes, faeces (manure, dung, faecal pellets, guano, frass), as well as products secreted, excreted or exuded from organisms (e.g., extracellular polymers, nectar, root exudates and leachates, dissolved organic matter, extracellular matrix, mucilage).” This broad definition of detritus includes nectar and root exudates, and therefore we should classify rhizosphere microorganisms, nectar feeders (e.g. butterflies, bees), and any body surface-dwelling symbionts as detritivores or saprotrophs. At present in the SFWO, we classify *nectarivore* as a subclass of *herbivore* because the opposite would consequently require classifying numerous nectar-feeding insects as detritivores if the ontology was to be expanded to aboveground food webs.

Another topic of discussion was the nature of the *predation* concept. It was argued that predation is not characterised by the consumption of a particular type of food, but rather that it is a food acquisition strategy. Similarly to ECOCORE, from which we reuse the *predation* class, the SFWO does not make a distinction between the classification of organisms based on their food type (e.g. *insectivore*, *nematophage*, *herbivore*) or feeding strategies (e.g. *predator*, *filter-feeder*, *grazer*). Besides, the term *predator* is very commonly used in the soil literature to refer to animals or protists feeding on animal/protist prey. We decided to retain this widely accepted usage. However, this may be the subject of further discussion, which could include other such terms, e.g., *parasite*.

A final example of a non-trivial terminological (and ontological) issue concerns the place of *saproxylophagy* in the hierarchy of concepts. Saproxylophagy is the process of feeding on dead or decaying wood, and as such, it is a specialisation (subclass) of *xylophagy*, which describes the feeding activities of herbivorous animals whose diet consists primarily of wood (living or dead). However, this would lead to the conclusion that saproxylophagy is a form of herbivory, when in fact it is most commonly considered to be a form of detritivory. We have chosen to make *saproxylophage* a subclass of *phytosaprophage*, itself a subclass of *detritivore*, to reflect this common use of the term in the literature.

## Applications of the Soil Food Web Ontology

### Standardisation of trophic trait data using the SFWO

Like many other fields in ecology, soil ecology has seen a strong increase in the availability of data over the last few decades, including molecular, taxonomic, distributional, morphological measurements and functional trait data for all the major groups that make up soil biodiversity. This opens exciting opportunities for a holistic description of the soil biota. However, the **<u>heterogeneity</u>** of data arising from different research contexts, sometimes from projects with scanty resources for data curation and sharing, make the task of data compilation tedious [46]. Integrative trait-based research, in particular, is hampered by a lack of standardisation in data structure, taxonomic resolution, trait names, values and units, and metadata quality (e.g. geolocation, time and space of sampling, measurement method). The need for a core terminology or data standard has been identified as an urgent priority for accelerating trait-based science [47].

From this perspective, the development of the Ecological Trait-data Standard (ETS) [13], a unified vocabulary for describing ecological trait datasets, is an important step towards improved interoperability and reuse of these datasets for future data aggregation and synthesis initiatives. The ETS requires that trait names (as well as categorical trait values and units) be linked to unambiguous trait definitions in a published ontology if available. Yet, ontologies for traits are scarce. The SFWO is making a significant contribution to filling this gap in the field of soil trophic ecology, by providing the ontological resources needed to standardise trophic trait datasets. Figure 2 shows how such a dataset, namely the BETSI database of soil invertebrate traits [14], can be formatted as per the ETS by mapping trait names and values to concepts in the SFWO. Figure 2a is an excerpt of the BETSI dataset in its original format. In Figure 2b, trait names/values are linked to corresponding concepts in the SFWO using their unique identifiers (IRIs). This process, called **<u>semantic annotation</u>**, requires scientists to spend time describing the content of their data, although it is greatly facilitated by online ontology browsers (e.g., Ontobee [38], AgroPortal [48]) and can be automated to a certain extent [49]. In return, semantic annotations provide semantically rich information about the data which ensures that their intended meaning is preserved in the long term, while improving data discoverability and easing the data integration process.

**Figure 2.**
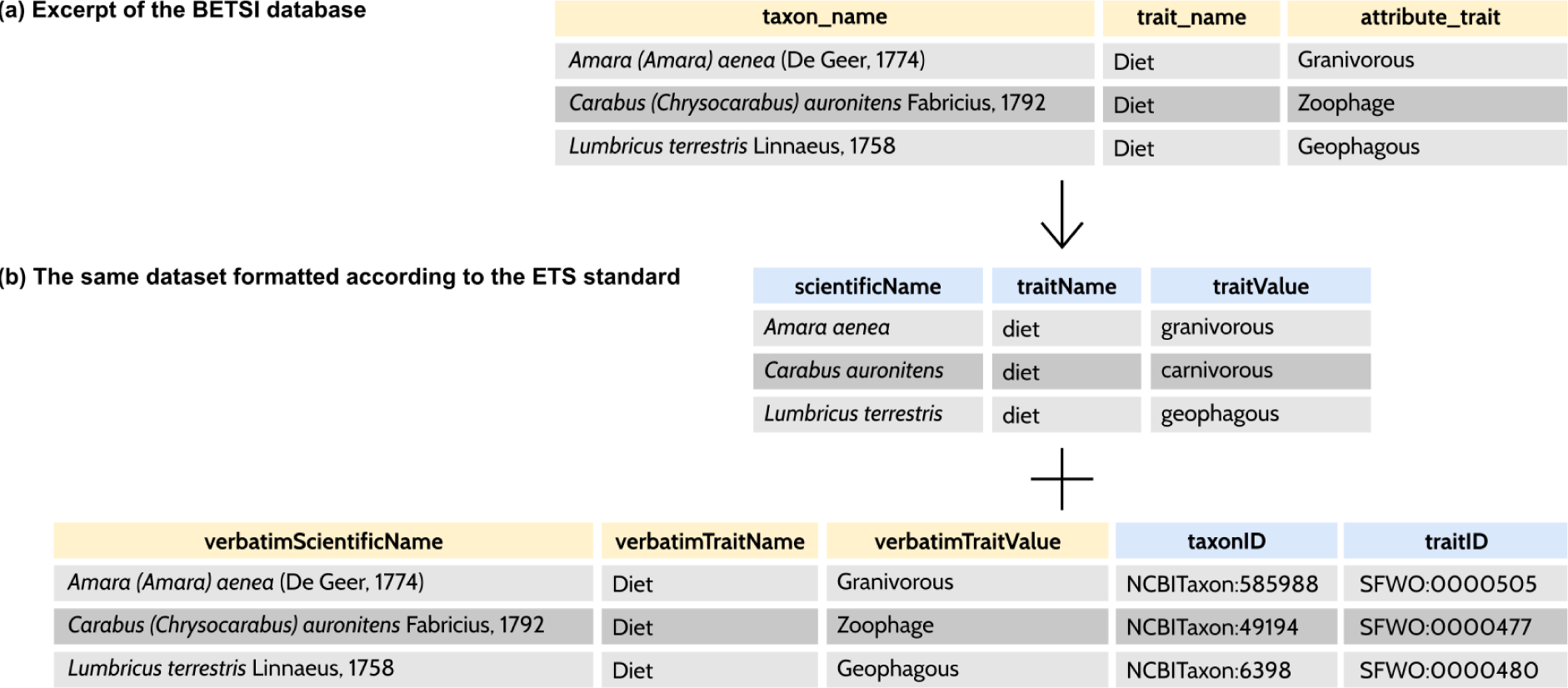
Standardisation of the BETSI database of soil invertebrate traits [14] using the ETS. (a) An excerpt from the BETSI database in its original format; BETSI uses terms from the T-SITA thesaurus for trait names and trait values. (b) The same dataset formatted according to the ETS: column names use terms from the ETS vocabulary; taxon scientific names, trait names, and trait values use terms from the NCBITaxon ontology and the SFWO; the IRIs of the corresponding ontology classes are added for taxa (taxonID) and traits (traitID).

### The SFWO as a mediator between heterogeneous data sources

Much ecological data reside in project-specific datasets, created by individual researchers or project teams to answer specific research questions. These datasets are typically small, focusing on a local set of taxa over relatively limited spatial and temporal scales, and formatted according to the needs of the project, often with no incentive to invest time and effort in data standardisation and transfer. Integrating these ‘long-tail data’ dispersed across different datasets could help address research questions at larger scales [46]. However, integrating ecological data is a difficult task because of the many dimensions of data heterogeneity that need to be addressed.

**<u>Semantic data integration</u>** aims to provide data users with the ability to seamlessly manipulate heterogeneous data from multiple sources by combining these data in a way that preserves their original meaning [50]. In practice, this often involves creating a mapping (i.e., semantic annotations) between the input data and an ontology that acts as a mediator to reconcile the semantic (i.e., terminological) heterogeneities between data sources. Schematic (i.e., structural) heterogeneities are then resolved by transforming the datasets into a common format and combining the transformed datasets into a single database [51].

Recently, graph-structured databases have become a popular choice for storing integrated data. **<u>Knowledge graphs</u>** store both the data and their semantic context (meaning) in the form of interconnected entities. This enables both humans and machines to better interpret and interact with the data. Knowledge graphs are flexible, easily accommodating new types of data, and efficient at handling large amounts of relational information. Knowledge graphs also support various inference and reasoning tasks, enabling the prediction of new relationships, the detection of inconsistencies and the validation of existing knowledge [52, 53].

Knowledge graphs have proven useful in many different domains, including biomedicine [54] and geoscience [55]. Soil food-web research would benefit greatly from a knowledge graph integrating trophic information from existing databases covering different soil taxonomic groups across several trophic levels and from the literature. This would provide a unified access to semantically-rich multigroup, multimorphic, multitrophic, and multisource information, while offering the ability to derive additional knowledge from the integrated data using automated reasoning or **<u>relational machine</u> <u>learning</u>**. Figure 3 shows how the SFWO can be used to combine the BETSI database with observed and well-documented trophic interactions from the Global Biotic Interactions (GloBI) database into a trophic knowledge graph using, for instance, the knowledge graph construction approach from Le Guillarme and Thuiller [21]. The resulting knowledge graph provides a single access point to harmonised and consolidated multisource data, thus greatly facilitating integrative studies across taxonomic groups and/or trophic levels.

**Figure 3.**
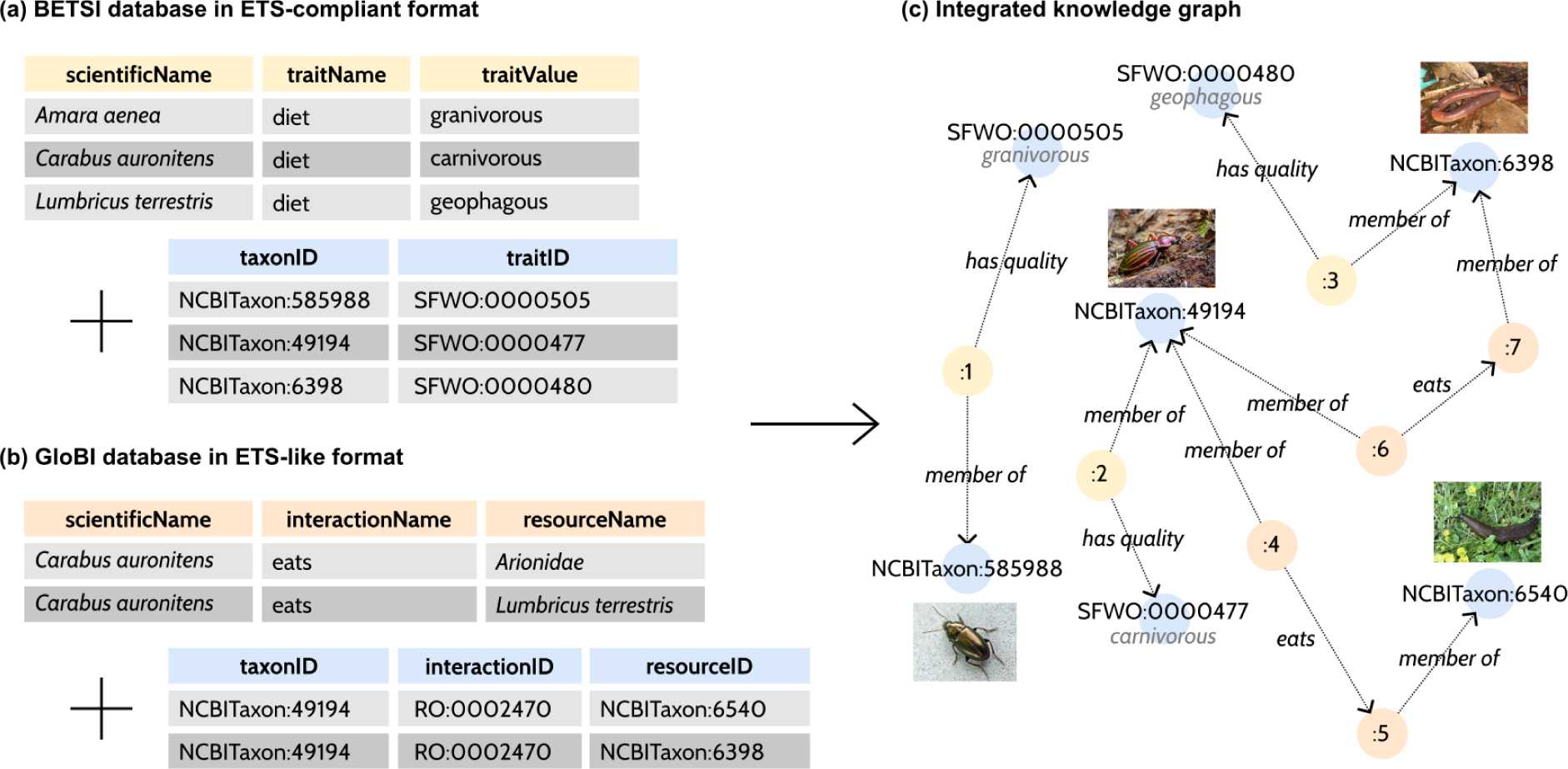
Semantic data integration provides a unified view over multisource data. Here, this unified view takes the form of a knowledge graph, a graph-structured knowledge base. (a) An excerpt from the BETSI database formatted according to the ETS. (b) An excerpt from the GloBI database transformed into an ETS-like format (as the ETS does not provide terms for interaction data). Semantic heterogeneities are resolved by linking the input data to the corresponding concepts in a target ontology or set of ontologies, here the SFWO, the Relations Ontology and the NCBITaxon ontology. (c) Semantically-annotated data are transformed into a graph format and combined into a single knowledge graph. The colour of a node represents the original source of the data or concept: BETSI (yellow), GloBI (orange), or one of the target ontologies (blue).

### Ontology-based information extraction and retrieval with the SFWO

Understanding soil food web structure at larger scales through integrative studies relies on our ability to compile data on the taxonomy, morphology, and trophic ecology of soil organisms from previously published research. Although nowadays the publication of datasets alongside scientific articles or data papers is greatly encouraged by publishers and facilitated by file hosting services such as Zenodo and Data Dryad, much ecological knowledge is still only available in human-readable form in the texts of journal articles, books, technical reports, grey literature and websites. In addition, the volume of scholarly texts is growing exponentially, making it even more difficult to find the relevant information and translate it into machine-readable form [56].

Computational approaches can help to make this valuable content available for large-scale ecological studies. This includes information retrieval algorithms that suggest documents of potential interest to the reader, and information extraction methods for transforming raw text into structured datasets [25]. Ontologies are valuable resources for information retrieval. For instance, ontology-based literature search is a feature of the Specialised Information Service Biodiversity Research (BIOfid) portal, which provides easy access to current and historical biodiversity literature [23]. Within the BIOfid portal, biodiversity texts are first semantically annotated with concepts from a reference ontology and indexed for efficient search. At search time, the BIOfid portal determines which ontological concepts are contained in the user query and presents the user with all relevant documents, including documents that do not necessarily contain the search terms *verbatim*, but may contain a synonym or a term referring to a sub-concept. Ontologies can also play a crucial role in the information extraction process [24]. As information extraction aims at retrieving specific information related to a particular domain of interest, a formal and explicit specification of the concepts of this domain by means of an ontology can be very useful in guiding the extraction process.

The SFWO has the potential to support both information retrieval and information extraction (Figure 4). By considering the SFWO as a dictionary containing terms of interest to soil trophic ecology (e.g., diets, food resources, keywords describing trophic interactions), one can use a **<u>dictionary-based</u> <u>Named Entity Recognition</u>** (NER) engine to locate mentions of these terms in digitised texts (Figure 4a). This process can be used to annotate documents for subsequent searching (Figure 4b). Incorporating trophic concepts from the SFWO into a semantic document retrieval engine could help soil ecologists screen the literature and more efficiently identify articles containing information relevant to soil food-web modelling and analysis. NER can also be used as a first step in extracting relational information from the literature (Figure 4c). Existing approaches have used the frequency of term co-occurrences to discover ant-plant associations [57], or “trigger words” to discover microorganism-habitat and microorganism-phenotype associations [58]. In combination with a taxonomic NER system able to detect mentions of taxa in text [59], the SFWO makes it possible to use similar approaches to discover taxon-diet or taxon-resource associations within a corpus of textual documents, and use the extracted information to populate literature-based datasets.

**Figure 4.**
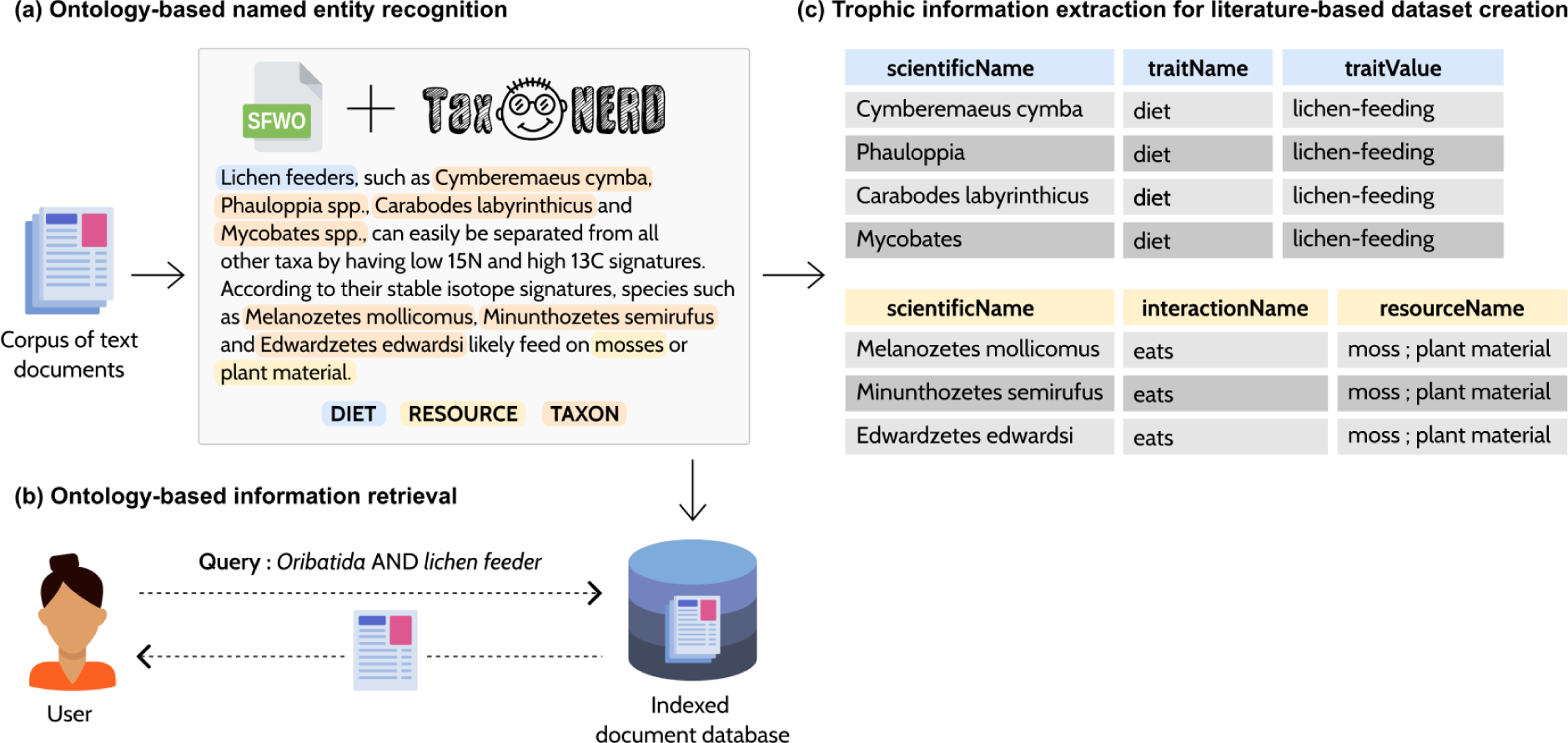
The SFWO can help scientists to access information on the trophic ecology of soil organisms that is scattered in the literature. (a) Named entity recognition (NER) is used to identify occurrences of terms from the SFWO into text documents. Similarly, a taxonomic NER tool like TaxoNERD [59] can locate occurrences of taxon mentions in text. (b) Annotated documents are indexed and can be efficiently retrieved using semantic queries. (c) NER is also a prerequisite for information extraction, which aims at building structured datasets from the literature.

### Applying automated reasoning to soil food-web research

Soil food-web research deals with a diversity of concepts (trophic interactions, trophic processes, diets, trophic groups, food resources, taxa) describing the trophic ecology of soil organisms. These concepts are tightly linked to each other by logical relationships. For instance, the diet (e.g., malacophagous) of an organism can be deduced from the trophic interactions it is involved in (e.g., feeding on molluscs). Similarly, organisms can be assigned to taxo-trophic groups (as per the classification by Potapov et al. [4]) provided that their taxonomic and trophic classifications are known. For example, knowing that *Carabus auronitens* is a malacophagous carabid, it is possible to assign this species to the group *Coleoptera.predators.Carabidae* (Figure 5). The SFWO uses axiomatized class definitions to encode these logical relationships between trophic concepts. This makes it possible to employ automated reasoners to evaluate the logical implications of the knowledge encoded in the ontology on explicitly stated data. This ability to apply reasoning to trophic data can support soil food-web research in several ways, including (1) testing the consistency of food-web models, (2) enabling the comparison among soil food webs across the literature, (3) inferring missing information in trophic datasets — e.g., inferring missing information about an organism’s diet from trophic interaction data, and (4) automatically classifying soil consumers into standard trophic groups for food-web reconstructions. As can be seen from these examples, the SFWO helps to address some issues related to the incompleteness or inaccuracy of trophic trait datasets, the heterogeneity in the resolution of food webs constructed using inconsistent trophic group definitions, or the burden of manual trophic group assignment.

**Figure 5.**
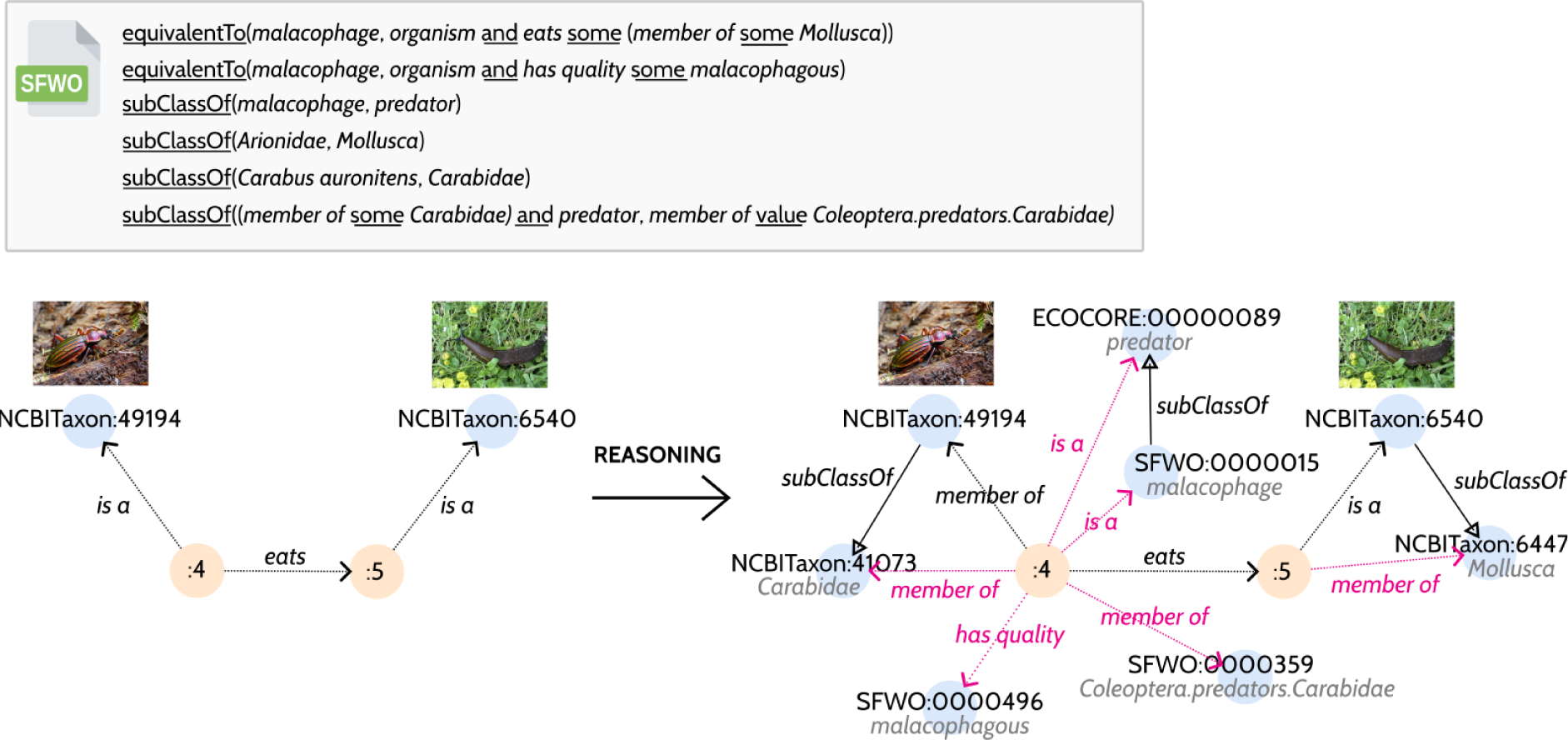
The SFWO provides class axiomatizations that can be used as logical rules to derive additional (implicit) knowledge from the input data using a reasoner. The graph on the left is an excerpt of the knowledge graph from *Figure 3* representing the (explicitly-stated) information about a trophic interaction between Carabus auronitens and Arionidae. A reasoner can combine this information with the knowledge encoded in the SFWO (namely that a malacophage is an organism that feeds on molluscs) and the NCBITaxon ontology (Arionidae is a subclass of Mollusca, C. auronitens is a subclass of Carabidae) to deduce that C. auronitens is a malacophage (possessing the quality malacophagous), therefore a predator and a member of the trophic group Coleoptera.predators.Carabidae. The graph on the right shows some of the additional facts that are derived by the reasoner (pink arrows).

## Conclusions and way forward

The SFWO is a collaborative and ongoing effort at developing a standard conceptual model of soil trophic ecology that is both interpretable by humans and computers. Building on existing ontologies, continuously curated and enriched by soil scientists in constant dialogue with an ontology engineer, the SFWO is an unprecedented resource for better knowledge management in the field. The SFWO helps to resolve terminological ambiguities and facilitates knowledge exchange between experts from different subfields by providing agreed definitions for over 500 concepts relevant to soil food-web research. It provides a formal conceptualization of the field of soil trophic ecology as a set of interrelated and uniquely-identified classes that can be used to harmonise trophic datasets through semantic annotation, thus facilitating data integration and discovery. The SFWO is also a useful resource for making the vast amount of unstructured information currently available only in the literature easier to find and use as part of larger-scale ecological studies. Finally, the logical foundations of the OWL ontology language allow a reasoner to use the knowledge encoded in the SFWO to infer additional information from explicitly-stated data, e.g., infer an organism’s diet and trophic group from its trophic interactions. All this makes the SFWO an essential tool for helping soil scientists to collect, annotate, interpret, (re)use and share trophic data, while opening many avenues for ontology-based applications in soil trophic ecology, including trophic knowledge graph construction, computer-assisted food web reconstruction, trophic information extraction, etc.

In its current state, the SFWO is the most comprehensive conceptual framework (and vocabulary) for describing trophic knowledge related to soil biota. However, the SFWO is likely missing some trophic concepts and many synonyms. In the next development iterations, we plan to add more synonyms, with the increased effort of adding them in multiple languages. We will pursue the axiomatization of trophic group classes, the definition of some of them involving non-trophic traits. For instance, the group *Lumbricina.all* is subdivided into three subgroups: *Lumbricina.epigeic, Lumbricina.anecic, Lumbricina.endogeic* based on the vertical stratification of their microhabitat. Providing logical definitions for these classes will require incorporating the concepts of epigeic, anecic, and endogeic organisms which are not trophic concepts *per se*, although closely related to feeding behaviours. We also plan to refine and enrich the classification of trophic groups, particularly with regard to bacteria, which are currently absent. In the longer term, we aim to increase the comprehensiveness of the SFWO so that it becomes easier to account for the plasticity of feeding behaviour in certain taxonomic groups. A first step in this direction was the introduction of life stage concepts in the ontology. We would like to explore the possibility of introducing representational mechanisms that would help quantify uncertainty in diet assignment, resource preferences, and the strength of trophic interactions. Perhaps the most difficult aspect of developing an ontology is getting it widely adopted by the scientific community. A few factors can facilitate engagement of the community of soil scientists. First, the ontology must be consensual, which means that it must capture the knowledge of the domain in a way that is accepted by the majority, if not by all. To help build and maintain consensus over the years, we wanted the development of the SFWO to be a continuous collaborative process in which everybody can participate, along the lines of open source software development. Therefore, on behalf of the SFWO working group (the current composition of which can be found on the project’s website^10^), we invite soil scientists, data providers, and anyone interested in using the SFWO to contribute to the improvement of the ontology. Contributions can be thoughtful comments on open issues, requests for new terms, changes to existing contents (modifications, clarifications), etc. Requests should be submitted via the project’s issue tracker^11^. Creating a new issue in the tracker allows the SFWO editors and user community to discuss proposed changes, fixes, or additions. The members of the SFWO working group will meet regularly to vet and approve changes to the ontology. To facilitate the adoption of the SFWO and the dialogue between ontology developers and domain experts (likely unfamiliar with the arcana of ontology languages such as OWL), we have made the ontology available in more user-friendly formats. A browsable version of the SFWO is available online on AgroPortal^12^. It includes a graphical representation of the concepts hierarchy, accessible from the Visualization tab in the Classes view. In addition, each new/updated release of the SFWO will be accompanied by an export of the ontology in a tabular format (CSV file). The export file of the version of the SFWO accompanying this paper is provided as Supplementary Material. Finally, we are in the process of writing comprehensive documentation to help users better understand the architecture of the ontology and the design choices that led to that architecture. Examples and tutorials on how to use the SFWO for ontology-based applications will also be included in the documentation (soon accessible from the project’s website).

Eisenhauer et al. [60] identified better data management, including data standardisation and integration, as one of the prominent research frontiers in soil ecology. This requires data standards (from the less expressive controlled vocabularies to the more expressive ontologies) to aid data harmonisation and compilation. The SFWO aims to tackle this issue and provide the semantic resources needed to enable researchers to make best use of the existing and future data on soil trophic ecology. In addition, the SFWO opens avenues for developing practical ontology-based applications in the field. We envisage that this work will foster other ontology development initiatives addressing other aspects of soil ecology.

## Footnotes

1 http://agroportal.lirmm.fr/ontologies/SFWO

2 https://github.com/oborel/obo-relations

3 https://bioportal.bioontology.org/ontologies/ECOCORE

4 https://obofoundry.org/ontology/ncbitaxon.html

5 https://jfaldanam.gitlab.io/EUTaxO/

6 https://github.com/obophenotype/fungal-anatomy-ontology

7 https://github.com/oborel/obo-relations/issues/647

8 https://protege.stanford.edu/

9 https://github.com/soilfoodwebontology/sfwo

10 https://soilfoodwebontology.github.io/

11 https://github.com/soilfoodwebontology/sfwo/issues

12 http://agroportal.lirmm.fr/ontologies/SFWO

## Box 1. Glossary

Automated reasoning: the process of automatically computing logical inferences.
Axiomatization: the process of providing logical definitions of ontology classes in the form of logical axioms, i.e. logical statements that describe the relationships between concepts.
Class: the description of a concept in an ontology. Classes in an ontology are organised into a taxonomy using the subsumption (*subClassOf*) relation.
Data heterogeneity: differences in semantics (terminologies, meaning, interpretation), schema (data structures, formats), and syntax (models, languages) between data from possibly different data sources.
Knowledge graph: a knowledge base that uses a graph-structured data model to integrate data.
Named entity recognition: the task of automatically detecting and classifying mentions of entities of interest (e.g., taxa, food resources, diets) in text.
OWL ontology: a formal representation of concepts within a domain of interest (e.g., ecology) and relationships between those concepts, encoded using a formal ontology language such as the Web Ontology Language (OWL). OWL is the World Wide Web Consortium’s (W3C) standard for authoring ontologies. It is based on a subset of first-order logic.
Property: the description of a relation in an ontology. This can be a relation between two classes (*object property*) or between a class and a data type value (*datatype property*). Properties in an ontology are organised into their own taxonomy using the *subPropertyOf* relation.
Reasoner: or reasoning engine, a piece of software that can perform automated reasoning.
Relational machine learning: a subdiscipline of artificial intelligence and machine learning that is concerned with the statistical analysis of relational (graph-structured) data.
Semantic annotation: the process of linking data to classes in an ontology.
Semantic data integration: the process of combining data from different sources into a single, unified view using ontologies.
Top-level ontology: or upper ontology, an ontology which consists of very general terms that are common across all domains. Upper ontologies support interoperability among a large number of domain-specific ontologies by providing a common starting point for the formulation of definitions.

## Box 2. Abbreviations

BETSI: Biological and Ecological Traits of Soil Invertebrates
BFO: Basic Formal Ontology
BIOfid: Specialised Information Service Biodiversity Research
GloBI: Global Biotic Interactions
IRI: Internationalized Resource Identifier
NCBI: National Center for Biotechnology Information
NER: Named Entity Recognition
OWL: Web Ontology Language
OBO: Open Biomedical Ontologies
PATO: Phenotype And Trait Ontology
PCO: Population and Community Ontology
RO: Relations Ontology
SFWO: Soil Food Web Ontology
T: SITA-Thesaurus for Soil Invertebrate Trait-based Approaches

## Acknowledgements

The research was funded by the French Agence Nationale de la Recherche (ANR) through the GlobNet (ANR-16-CE02-0009) project, from the two ‘Investissement d’Avenir’ grants (MIAI@Grenoble Alpes: ANR-19-P3IA-0003, OSUG@2020: ANR-10-LAB-56), and the METRO Grenoble Alpes, the Departement Isere and the Office Français de la Biodiversité to Orchamp. Ideas also matured thanks to working group meetings of the EUdaphobase COST Action (CA18237) and sDiv funded sOilFauna project (grant SFW9.02). A.P. acknowledges support of the DFG Emmy Noether program (Project number 493345801) and of iDiv (DFG–FZT 118, 202548816). C.A.M.M. acknowledges the DFG grant MO 412/54-2, project number 326061700, supporting the Specialised Information Service Biodiversity Research (BIOfid, www.biofid.de).

## Author contributions

**Nicolas Le Guillarme:** Methodology, Software, Writing - Original Draft, Writing - Review & Editing **Mickael Hedde:** Conceptualization, Validation, Writing - Review & Editing, Supervision **Anton M. Potapov:** Conceptualization, Validation, Writing - Original Draft; Writing - Review & Editing **Carlos A. Martínez-Muñoz:** Conceptualization, Validation, Writing - Review & Editing **Matty P. Berg:** Validation, Writing - Review & Editing **Maria J.I. Briones:** Validation, Writing - Review & Editing **Irene Calderón-Sanou:** Validation, Writing - Review & Editing **Florine Degrune:** Validation, Writing - Review & Editing **Karin Hohberg:** Validation, Writing - Review & Editing **Camille Martinez-Almoyna:** Validation, Writing - Review & Editing **Benjamin Pey:** Validation, Writing - Review & Editing **David J. Russell:** Validation, Writing - Review & Editing **Wilfried Thuiller:** Writing - Review & Editing, Funding acquisition.

